# Absence of testes at puberty impacts functional development of nigrostriatal but not mesoaccumbal dopamine terminals in a wild-derived mouse

**DOI:** 10.1101/2025.05.16.654596

**Authors:** Jaewan Mun, Samantha Jackson, George Prounis, Chayarndorn Phumsatitpong, Niloofar Motahari, Lance Kriegsfeld, Markita Landry, Linda Wilbrecht

## Abstract

The nigrostriatal and mesoaccumbal dopamine systems are thought to contribute to changes in behavior and learning during adolescence, yet it is unclear how the rise in gonadal hormones at puberty impacts the function of these systems. We studied the impact of prepubertal gonadectomy on evoked dopamine release in male *Mus spicilegus*, a mouse whose adolescent life history has been carefully characterized in the wild and laboratory. To examine how puberty impacts the dopamine systems in *M. spicilegus* males, we removed the gonads prepubertally at P25 and then examined evoked dopamine release in the dorsomedial, dorsolateral, and nucleus accumbens core regions of striatal slices at P60-70. To measure dopamine release, we used near-infrared catecholamine nanosensors (nIRCats) to enable study of spatial distribution of dopamine release sites in each striatal region. We found that prepubertal gonadectomy led to a significantly reduced density of dopamine release sites and reduced dopamine release at each site in the dorsolateral nigrostriatal system compared to sham controls. By contrast, mesoaccumbal dopamine release was comparable between sham and gonadectomized groups. Our data suggest that during adolescence the development of the nigrostriatal dopamine system is significantly affected by the rise in gonadal hormones in males, while the mesoaccumbal system shows no detectable sensitivity at this time point. These data are consistent with molecular studies in rodents that suggest nigrostriatal neurons are sensitive to androgens at puberty, and extend our understanding of how gonadal hormones could impact the spatial distribution and release potential of dopamine terminals in the striatum.

**Significance Statement:** Here we use a wild-derived species, *Mus spicilegus*, to study adolescent development. This wild-derived species has value over standard lab mice because it is more likely to exhibit evolved developmental programs relevant to dispersal and other natural behaviors. By using this wild-derived species and metrics of evoked dopamine release with spatial resolution, we can test if the rise in gonadal hormones at puberty plays a role in maturation of dopamine terminal function in the striatum. These findings may help us better understand developmental programs in humans that orchestrate changes in behavior at adolescent milestones in contexts of both health and disease.

## Introduction

Across many species, adolescence is a time of exploration, sensation-seeking, and adaptive and sometimes maladaptive risk-taking (Dahl et al., 2018; Worthman and Trang, 2018; Natterson-Horowitz and Bowers, 2020; Lin and Wilbrecht, 2022). These behavioral changes are accompanied by significant structural and functional remodeling of the brain, including changes in dopamine neurons and their terminals in the striatum, (Larsen et al., 2020) as well as the neocortex (Hoops and Flores, 2017; Delevich et al., 2021). Changes in dopamine systems during adolescence are of particular interest because they may be implicated in changes in learning and decision-making as well as the development of mental health issues that emerge in adolescence (Ge et al., 2001; Graber et al., 2004; Deardorff et al., 2007; Mendle et al., 2018; Copeland et al., 2019; Simpson et al., 2022).

Here we focus on males and the development of dopamine in the nigrostriatal and mesoaccumbal pathways of the dopamine system. Several studies suggest nigrostriatal and/or mesoaccumbal dopamine neurons show dynamic changes across adolescent development in rodent brains. Multiple studies in rats show evoked dopamine release increases with development in males (Gazzara et al., 1986; Stamford, 1989; Pitts et al., 2020), but some show a decrease (Kuhn et al., 2010). Sex and regional differences can also differ, making consensus difficult.

Once rodents reach adulthood, tyrosine hydroxylase (TH) levels in nigrostriatal and mesoaccumbal neurons and striatal tissue are sensitive to activational effects of gonadal hormones, which can be isolated by examining the effects of gonadectomy (GDX) (i.e. castration) followed by hormone replacement (Abreu et al., 1988). The regulation of terminal release of dopamine in the striatum has also been found to be modulated by testosterone after GDX (Shemisa et al., 2006). However, not all results again agree or show a consistent direction of effect (Xiao and Becker, 1994). The effects of gonadal hormones on cells at puberty can have opposing effects in adulthood. For example, male rats that undergo GDX in adulthood exhibit a decrease in cocaine-stimulated dopamine “overflow” in the dorsal striatum, but rats that undergo GDX prior to puberty can exhibit an increase in dopamine “overflow” (Kuhn et al., 2010). In summary, there is a strong body of evidence suggesting gonadal hormones in males play a role in midbrain dopamine neuron development, but there are still open questions about the role of gonadal hormones at the time of puberty and their impact on terminal growth and function.

To clarify the role of gonadal hormones in dopamine neuron development, we studied the effect of prepubertal GDX (P25-28) on evoked dopamine release in the striatal dorsomedial (DMS), dorsolateral (DLS), and nucleus accumbens (NAc) regions in male *Mus spicilegus* during late adolescence (P60-70) using near infrared catecholamine sensors (nIRCats). This is a new imaging tool designed for measuring evoked catecholamine release that allows for spatial resolution of “hotspots” giving us more information than previously available about terminal function and density (Beyene et al., 2019). We opted to study a wild-derived species, *Mus spicilegus*, rather than standard laboratory mice, to identify ethologically relevant developmental programs that may link maturation of dopamine neurons to pubertal changes. *Mus spicilegus* is a particularly valuable species to study because their adolescent behavior has been documented in the wild (Garza et al., 1997; Gouat et al., 2003; Poteaux et al., 2008), and they show developmental changes in risk-taking and exploratory behavior in the lab (Groó et al., 2013; Lafaille and Féron, 2014; Cryns et al., 2022; Bárdos et al., 2024). We therefore expect if there are programs that couple dopamine system maturation to pubertal processes, that we will have 1) a high likelihood of detecting them in this wild-derived species, and 2) the significance of these behavioral changes may be interpretable in an ethological context.

## Materials and Methods

### Animals

Male *Mus spicilegus* were bred in-house. Mice were weaned on postnatal day (P)21 and group-housed on a 12:12h light:dark cycle. Animals were given access to food and water *ad libitum* and housed with nesting material. Mice underwent gonadectomy or sham surgery between P25 and P28. Between P60 and P70, mice were perfused and brains collected. All procedures were approved by the Animal Care and Use Committee of the University of California, Berkeley and conformed to principles outlined by the NIH Guide for the Care and Use of Laboratory Animals.

### Synthesis of nIRCats

HiPco SWCNTs and (GT)_6_ oligonucleotides were purchased from NanoIntegris, and Integrated DNA Technologies, respectively. 0.1 mM of (GT)_6_ and 1 mg of SWCNT were mixed in 1 mL of 100 mM NaCl solution. The mixture was first bath-sonicated (Branson Ultrasonic 1800) and then probe-tip sonicated (Cole-Parmer Ultrasonic Processor, 3-mm tip in diameter, 5 W power) in an ice water bath for 10 minutes each. The resulting suspension was centrifuged at 16,800g for 90 minutes to remove unsuspended SWCNT aggregates. The supernatant was collected for further purification. To remove excessive free (GT)_6_ in the recovered suspension, Amicon spin filter tubes (MW 100k) were used. 500 µL of the suspension was spin-filtered at 8,000g for 5 minutes, and the filtrate was discarded. 500 µL of molecular-grade water was added to the spin filter tube and was centrifuged at 8,000g for 5 minutes. This wash step with water was repeated three times. To recover concentrated (GT)_6_-SWCNT suspension, the filter tube was reversed into a new bottom container and spun at 1,000g for 5 minutes. The final sample was collected and used for characterization.

### Characterization of nIRCats

The concentration of the collected (GT)_6_-SWCNT was characterized by its absorbance at 632 nm, which was measured by Nanodrop (Thermo Scientific). An extinction coefficient of 0.036 L mg^−1^ cm^−1^ at 632 nm was used. Each batch of nIRCats was then diluted to 5 mg L^−1^ in 0.1 M NaCl for in vitro fluorescence measurements. 99 µL (GT)_6_-SWCNT was placed in a well of a 384-well plate. Fluorescence spectra were collected with an inverted Zeiss microscope (20x objective lens, Axio Observer D1) coupled to a spectrometer (SCT 320, Princeton Instruments) and a liquid nitrogen-cooled InGaAs detector (PyLoN-IR, Princeton Instruments). A 721 nm laser (Opto Engine LLC) was used as an excitation light source. To observe the response of nIRCats to dopamine, fluorescence was measured before and after 1 µL of 100X diluted dopamine hydrochloride (Sigma Aldrich) was added to the nanosensor. nIRCats increase in fluorescence by up to 3,500% in the presence of catecholamines

### Gonadectomies

To examine adolescent development in the absence of steroid sex hormones, gonadectomies (GDX) were performed before puberty onset between P25 and P28 as described previously (Delevich et al., 2020). Before surgery, mice were injected with analgesics 0.05 mg/kg buprenorphine and 10 mg/kg meloxicam and anesthetized with 1-2% isoflurane during surgery. The incision area was shaved and scrubbed with ethanol and betadine. Ophthalmic ointment was applied over the eyes to prevent drying. A 1 cm incision was made with a scalpel in the lower abdomen across the midline to access the abdominal cavity. The testes were clamped off from the spermatic cord with locking forceps and ligated with sterile sutures. After ligation, the gonads were excised with a scalpel. The muscle and skin layers were sutured, and wound clips were placed over the incision for 7–10 days to allow the incision to heal. An additional injection of 10 mg/kg meloxicam was given 12–24 hours after surgery. Sham control surgeries were identical to GDX except that the gonads were simply visualized and were not clamped, ligated, or excised. Mice were allowed to recover on a heating pad until ambulatory and were postsurgically monitored for 7–10 days to check for normal weight gain and signs of discomfort/distress. Mice were co-housed with 1-2 siblings who received the same surgical treatment. Subjects in this study included 6 intact and 6 castrated males.

### Acute brain slice preparation and nanosensor incubation

Acute brain slices were prepared from adult mice aged between P60 and P70 following previously reported protocols (Piekarski et al., 2017). Briefly, mice were deeply anesthetized via isoflurane exposure, followed by an overdose of ketamine/xylazine solution. Transcardial perfusion was performed by injecting ice-cold cutting buffer (119 mM NaCl, 26.2 mM NaHCO_3_, 2.5 mM KCl, 1 mM NaH_2_PO_4_, 3.5 mM MgCl_2_, 10 mM glucose, and 0 mM CaCl_2_). 300 µm thick coronal slices were cut in ice-cold cutting solution before being incubated in ACSF buffer (119 mM NaCl, 26.2 mM NaHCO_3_, 2.5 mM KCl, 1 mM NaH_2_PO_4_, 1.3 mM MgCl_2_, 10 mM glucose, and 2 mM CaCl_2_). Slices were bubbled with 95% O_2_/5% CO_2_ at 37 °C for 30 minutes, then incubated at room temperature. Slices were incubated in a small incubation chamber (Scientific Systems Design Inc.) with 2 mg L^−1^ nIRCats in 5 mL ACSF buffer bubbling with 95% O_2_/5% CO_2_ for 15 minutes. The slices were washed with ACSF buffer to remove unlabeled nIRCats and kept in ACSF buffer for another 15 minutes before imaging.

### nIRCat imaging in *ex vivo* slices using nIR microscopy

Dopamine imaging was performed *ex vivo* with a modified upright epifluorescent microscope. Details of the microscope are described in a previous study (Beyene et al., 2019). Excitation of nIRCats was done via a 785 nm laser, and fluorescence was measured with an InGaAs detector (Ninox 640). A slice labeled with nIRCats was transferred onto the sample stage of the microscope. The surface of the slice was made in contact with a bipolar stimulation electrode using 4x and 60x objective lenses. Specifically, the stimulator was positioned next to the imaging field of view in DMS, DLS, or NAc. All stimulation experiments were recorded with a frame rate of 9 frames per second and single-pulse electrical stimulations were applied after 200 frames of baseline were acquired. Each video acquisition lasted 600 frames. Within a field of view, stimulation amplitudes were repeated three times, with 5 minutes in between for terminal recovery.

### Data analysis of nIRCat fluorescence response

A python-based image analysis tool (https://pypi.org/project/NanoImgPro/0.2.3/) was used to process a raw 600-frame image stack. First, a field of view was divided into multiple regions (7 µm x 7 µm in size), and ROIs were identified when they showed a significant change in fluorescence over the background after electrical stimulation. Specifically, if the fluorescence change in a given region exceeded three times the standard deviation of baseline fluctuations, it was considered dopamine-induced. Regions that exhibited significant changes due to dopamine were defined as active release sites (active ROIs). Release site density was calculated by dividing the number of active ROIs by the total number of regions.

In each ROI, ΔF/F_0_ was achieved by dividing the change of fluorescence by the baseline. Next, time constants of signal onset (τ_on_) and decay (τ_off_) were extracted by approximating the ΔF/F_0_ vs time curve as a product of exponential decay functions as follows.

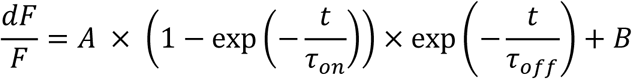

Here, A and B are constants. To compare ROI number, peak ΔF/F_0_, and τ_off_ between GDX and sham, mean values of the parameters were used. Each parameter was first averaged in a single field of view and further averaged over different fields of views within the same slice and across slices.

### Fecal testosterone collection and analysis

Fecal matter (0.4g) were collected from home cages of group-housed animals reared on a 12:12h light:dark cycle every 10 days from P20-P60. All mice were placed in new cages the day before collection to minimize steroid decay due to old fecal samples. Fecal samples were stored at −20°C until processed. Prior to testosterone extraction, fecal samples were dried and ground into a powder. Drying was performed by placing the samples in an oven at 65°C for a maximum of 90 minutes. The dried samples were then ground into a fine powder using a coffee grinder and transferred to 1.5 mL Eppendorf tubes using a funnel. The tubes were subsequently weighed to adjust the sample weight to 0.2 g, with precise weights recorded for all samples. Steroids were extracted following the Steroid Solid Extraction protocol from Arbor Assays. Briefly, 0.2g of dried fecal material was mixed with 1.8 mL of ethanol and shaken vigorously for 30 minutes. The samples were then concentrated by air dry in the oven at 65°C and re-suspended in 100 µL of ethanol. The concentrated extract was then diluted further with Assay Buffer. Fecal testosterone was determined in duplicate aliquots using the DetectX testosterone enzyme immunoassay kit (Arbor Assays, Ann Arbor, MI). The intra-assay coefficient of variation for this assay was 8.48%. Assay sensitivity was 9.92 pg/mL.

## Results

After weaning at P21, male *M. spicilegus* were gonadectomized at P25-28 or underwent sham surgery as a control (Figure 1a). Age at gonadectomy was chosen based on data from *Mus musculus* (Piekarski et al., 2017) and a preliminary study of fecal testosterone (T) of *M. spicilegus* in which we found a ∼ 50% rise in T between P30-90 (Figure 1a).

**Figure 1.**
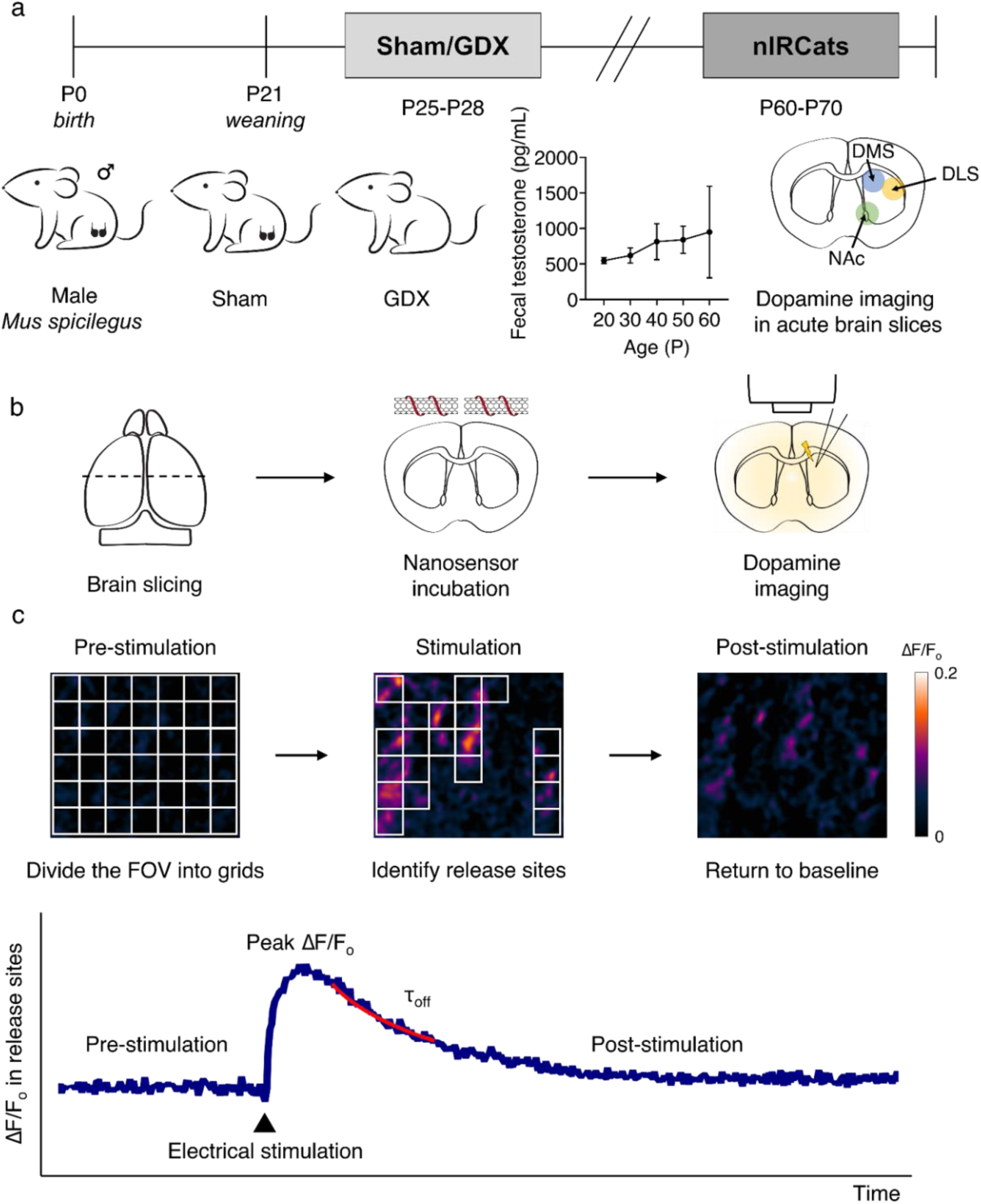
Schematic of the experimental timeline using nIRCat nanosensor based technology. (a) Male *Mus spicilegus* were gonadectomized (GDX) after weaning at P25-28 to remove gonads before onset of puberty. Data from a separate cohort of male *M. spicilegus* (n=4 cages) show an increase in average fecal testosterone from P20-60 in this species. Error bars represent the SEM. Dopamine imaging in acute brain slices from sham / GDX mice was performed during late adolescence or cusp of early adulthood (P60-P70) in this species reared on this 12h:12h light cycle. (b) Schematic of imaging experiments in brain slices. Acute brain slices were prepared and labeled with dopamine nanosensors. Electrically evoked dopamine release was monitored in near-infrared with a custom-built microscope. (c) Schematic of data analysis for nanosensor fluorescence response in brain slices. Three metrics of dopamine release (density of active ROIs as fraction of total ROIs, peak ΔF/F within active ROIs, and τ_off_) were extracted from the nIRCat fluorescence response and used for comparison between groups.

To examine the effect of GDX on the function of the nigrostriatal and mesoaccumbal dopamine system, we extracted the brains of both GDX and sham mice during late adolescence/early adulthood (P60-70) and acute brain slices were prepared following previously reported protocols (Beyene et al., 2019; Yang et al., 2021) (Figure 1b). Next, we imaged electrically-evoked nIRCat responses in three striatal regions: the DLS, the DMS, and the NAc using a custom-built microscope with a near-infrared (nIR) camera. Dopamine release was evoked using a 1 ms single electrical pulse (0.3 mA) from a bipolar stimulating electrode and ΔF/F_0_ was calculated for all pixels with nIRCat labeling. We divided each field of view (178 µm x 142 µm) into 546 regions of interest (ROIs) (7 µm x 7 µm in size) and measured the ΔF/F_0_ for each ROI (Figure 1c). We identified ROIs with significant fluorescence modulation following stimulation and designated them as dopamine release sites (active ROIs). Specifically, if the fluorescence change in a given region after stimulation exceeded three times the standard deviation of baseline fluctuations, it was considered evoked dopamine-induced. Since nIRCats fluorescence modulation is dopamine-dependent, the fraction of active ROIs serves as a proxy for dopamine release site density. Next, we analyzed the change of ΔF/F₀ at active release sites over time to characterize dopamine release decay dynamics after stimulation (Figure 1c).

### Prepubertal GDX males showed fewer and weaker active dopamine release sites in the dorsal striatum than sham controls

We found that a single pulse of electrical stimulation induced clear fluorescence modulation of nIRCats in the DLS (Figure 2a). The fluorescence of nIRCats returned to baseline following an exponential decay post-stimulation, enabling the use of the same field of view for subsequent stimulations (Figure 2a,b). Overall, GDX mice exhibited significantly smaller evoked dopamine release upon electrical stimulation in the DLS than the sham cohort. When comparing GDX and sham groups in metrics of density of active release sites (defined here as active 7×7 ROIs) and average peak ΔF/F_0_ in each release site, we found that GDX mice showed a significantly lower density of active release sites (expressed as a % of total sites) (Figure 2d) (p<0.05) and significantly smaller average peak ΔF/F_0_ in each of these release sites (active ROIs only) (Figure 2c) (p<0.05). Of course, dopamine released from terminals upon electrical stimulation is a function of both the density of terminals and release per terminal. Our results suggest that prepubertal GDX affected both the density of terminals, as measured by % density of active sites, and release per terminals, as measured by peak ΔF/F_0_ per active site in the DLS.

**Figure 2.**
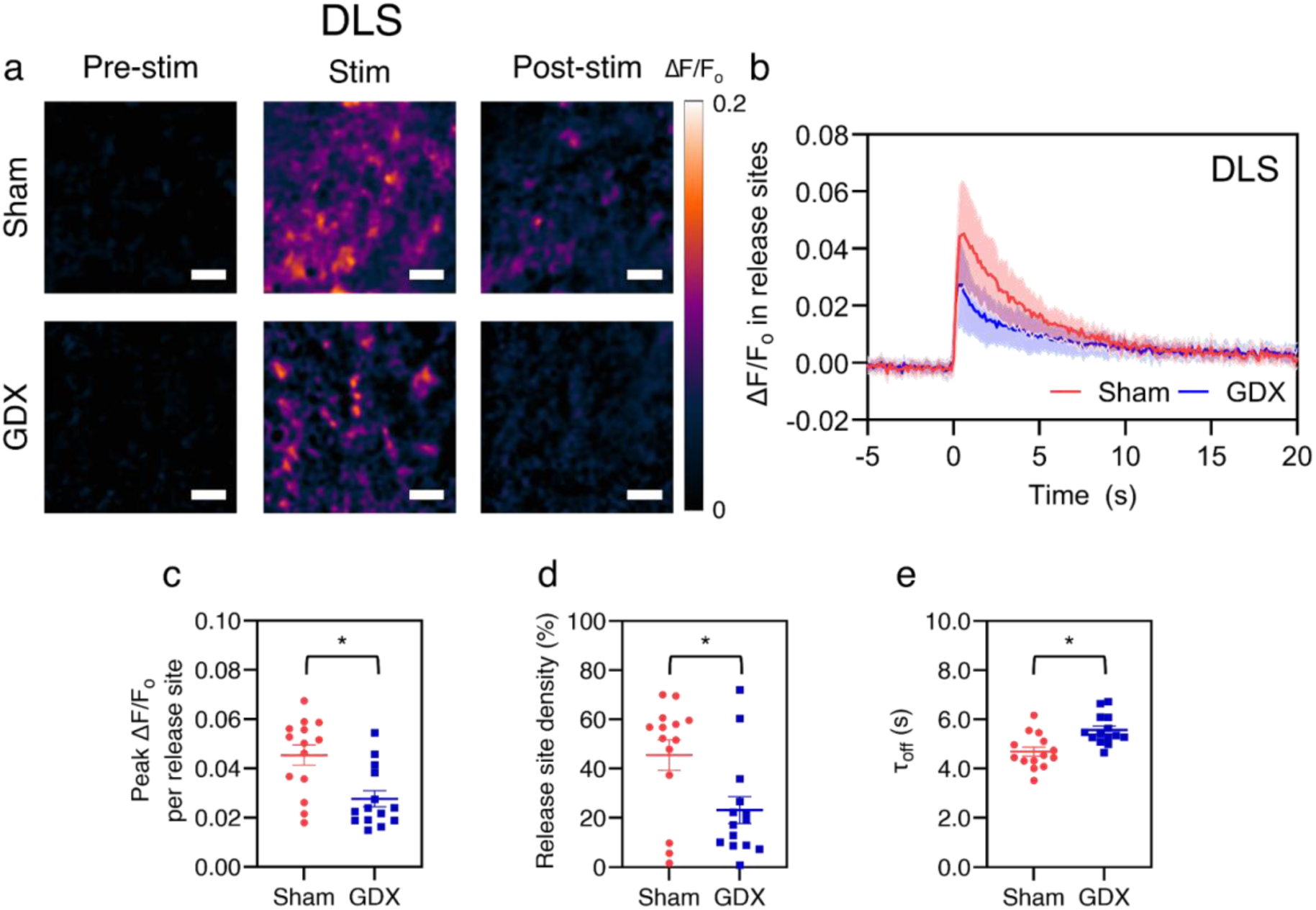
The density of dopamine release sites and release at each site was lower in the DLS of adult mice that underwent prepubertal GDX compared to sham controls. (a) Repeat images of the same field of view in the DLS and ΔF/F_0_ of nIRCat signal in sham (top) and GDX (bottom) mice. “Pre-stim” is before electrical stimulation is applied, “Stim” is the peak ΔF/F_0_ following stimulation, and “Post-stim” is when fluorescence is returning to baseline. (b) Average change in ΔF/F_0_ in the DLS in response to 0.3 mA electrical stimulation applied at 0 s in sham (n=14 brain slices/6 mice) and GDX (n=14 brain slices/6 mice) mice. Change in fluorescence reflects electrically-evoked dopamine release at time 0. Shaded area represents standard deviation. Metrics of dopamine release based on nIRCat signal per slice are summarized in (c)-(e). (c) Average peak ΔF/F_0_ at active release sites (defined as 7 µm squares regions of interest (ROIs) where release was at least three times the standard deviation of baseline fluctuation after electrical stimulation). (d) Active release site density (% of total sites in FOV that were defined as active), and (e) τ_off_ calculated using ΔF/F_0_ from only active release sites (active 7×7 µm ROIs). Scale bars, 10 µm. *p<0.05. Error bars represent the SEM.

Next, we compared the decay of the evoked signal which may reflect dopamine reuptake dynamics. We quantified this reuptake feature using an exponential decay function (Figure 1c, 2b). The τ_off_ was extracted from the curve to measure the decay time from peak fluorescence back to baseline. We found that GDX mice exhibited larger τ_off_ (Figure 2e)(p<0.05), suggesting that a lack of gonadal hormones at puberty may also block an increase in dopamine reuptake in the DLS seen in adulthood.

We next investigated the DMS, as it is well known that the DLS and DMS, while both targets of nigrostriatal dopamine play different roles in behavior (Yin and Knowlton, 2006). In the DMS, as in the DLS, the GDX group showed significantly reduced peak ΔF/F_0_ per active release site compared to the sham group (p<0.05)(Figure 3a-c). However, in the DMS, no significant difference was observed in the density of active release sites (Figure 3d). This result suggests that in the DMS, the pubertal rise in gonadal hormones in males may not significantly affect the number of dopamine release sites, but does influence the amount of dopamine released per release site. In the DMS, we found no significant difference in τ_off_, suggesting dopamine reuptake in DMS was not sensitive to gonadal hormones at puberty (Figure 3e).

**Figure 3.**
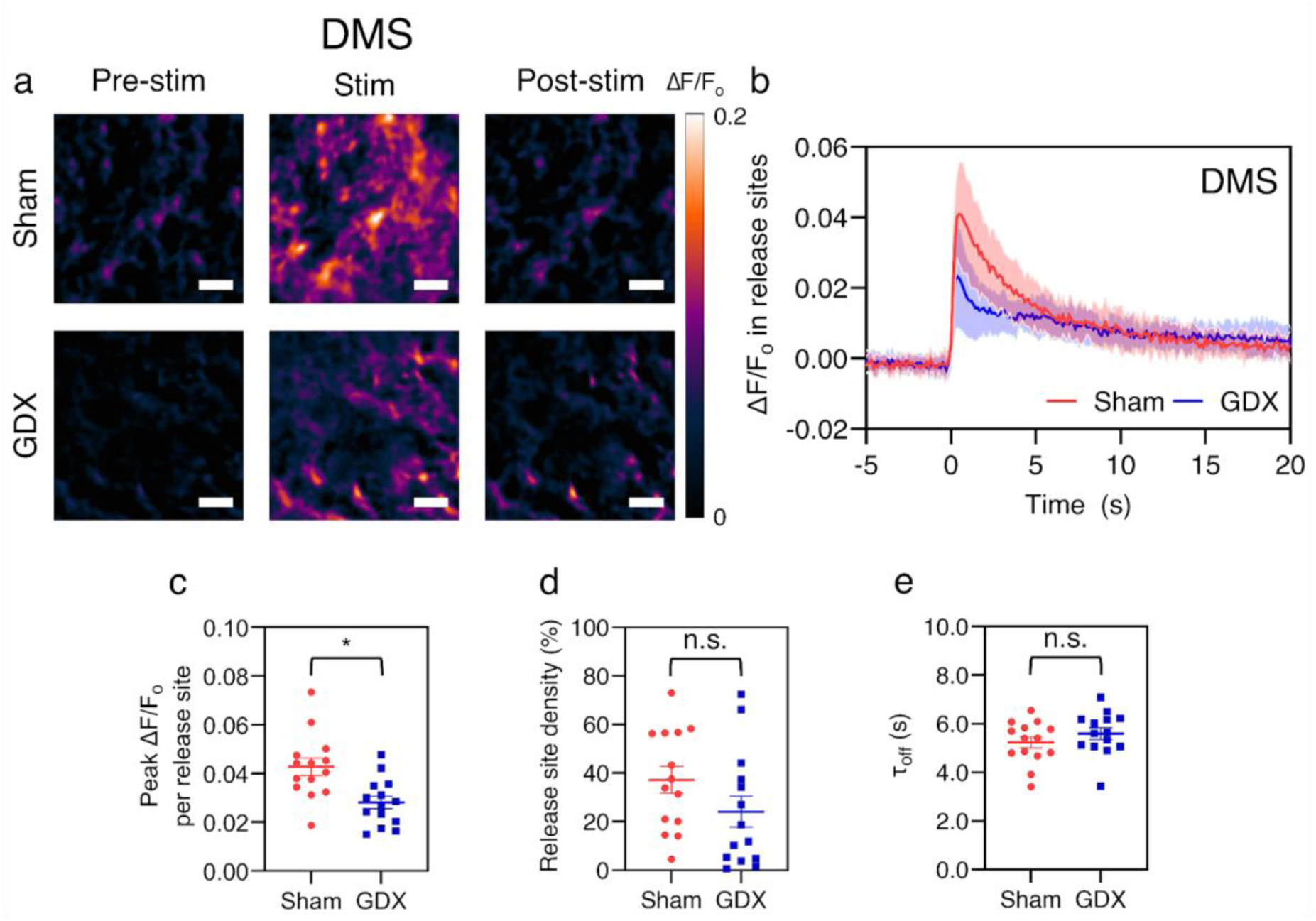
Dopamine release at active sites was lower in the DMS of adult mice that underwent prepubertal GDX compared to sham controls. (a) Repeat images of the same field of view in the DMS and ΔF/F_0_ of nIRCat signal in sham (top) and GDX (bottom) mice. “Pre-stim” is before electrical stimulation is applied, “Stim” is the peak ΔF/F_0_ following stimulation, and “Post-stim” is when fluorescence is returning to baseline. (b) ΔF/F_0_ change in DMS in response to 0.3 mA electrical stimulation applied at 0 s in sham (n=14 brain slices/6 mice) and GDX mice (n=14 brain slices/6 mice). Change in fluorescence reflects electrically-evoked dopamine release at time 0. Shaded area represents standard deviation. Metrics of dopamine release based on nIRCat signal per slice are summarized in (c)-(e). (c) Average peak ΔF/F_0_ at active release sites (defined as 7×7 µm square regions of interest (ROIs) where release was at least three times the standard deviation of baseline fluctuation after electrical stimulation). (d) Active release site density (% of total FOV defined as active), and (e) τ_off_ calculated using ΔF/F_0_ from only active release sites (active 7 µm ROIs). Scale bars, 10 µm. *p<0.05. Error bars represent the SEM.

### Prepubertal GDX had little impact on evoked dopamine release in the ventral striatum

Finally, we examined dopamine release in the NAc core in the same mice used above. Dopaminergic projections to the ventral striatum (VS), including the NAc, originate from the ventral tegmental area in the midbrain. This projection is also called the mesoaccumbal dopamine pathway. We found that GDX and sham mice showed comparable metrics of evoked dopamine release when quantified for the whole field of view (Figure 4b), ΔF/F_0_ per active release site (Figure 4c), and density of release sites (Figure 4d). The τ_off_ decay time constants were also comparable (Figure 4e). These data suggest that gonadal hormones at puberty in male *M. spicilegus* mice had little impact on the mesoaccumbal dopamine system terminals.

**Figure 4.**
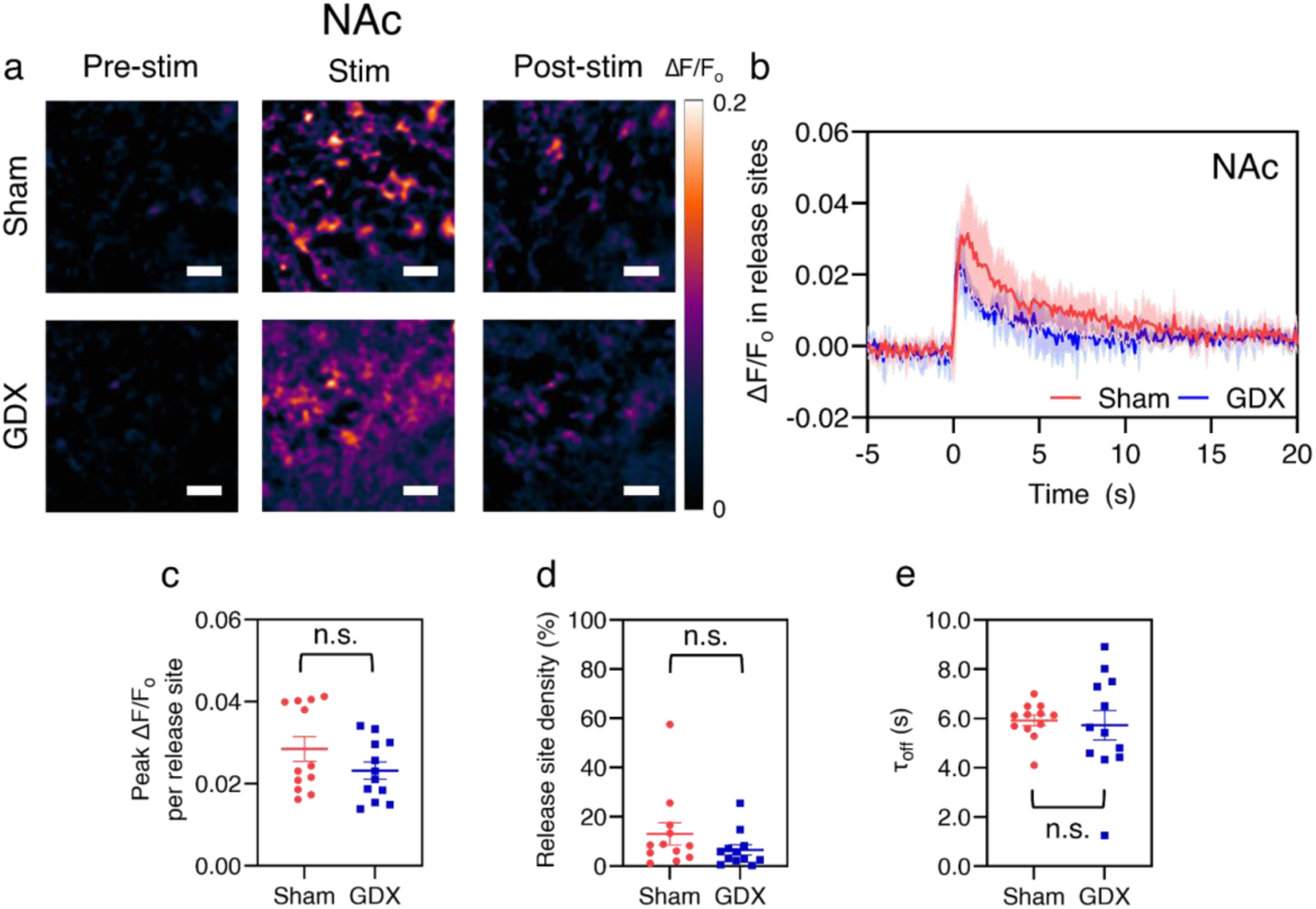
Evoked dopamine in the NAc at P60 was not significantly affected by prepubertal GDX in *M. spicilegus.* (a) Repeat images of a field of view in NAc and ΔF/F_0_ of nIRCat signal before, following, and after electrical stimulation in sham (top) and GDX (bottom) mice. “Pre-stim” is before electrical stimulation is applied, “Stim” is the peak ΔF/F_0_ following stimulation, and “Post-stim” is when fluorescence is returning to baseline. (b) ΔF/F_0_ change in NAc in response to 0.3 mA electrical stimulation applied at 0 s in sham (n=12 brain slices/5 mice) and GDX mice (n=12 brain slices/5 mice). Change in fluorescence reflects electrically-evoked dopamine release at time 0. Shaded area represents standard deviation. Metrics of dopamine release based on nIRCat signal in each slice are summarized in (c)-(e). (c) Average peak ΔF/F_0_ at active release sites (defined as 7 µm squares regions of interest (ROIs) where release was at least three times the standard deviation of baseline fluctuation after electrical stimulation). (d) Active release site density (% of total FOV defined as active), and (e) τ_off_ calculated using ΔF/F_0_ from only active release sites (active 7 µm ROIs). Scale bars, 10 µm. *p<0.05. Error bars represent the SEM.

To study similarities and differences between striatal regions, we also plotted metrics of dopamine release side-by-side in a new panel. We then compared metrics of dopamine release across the DMS, DLS, and NAc using ANOVA followed by post-hoc analysis with Bonferroni correction. We found that, in sham males, dopamine release was significantly greater in the dorsal striatum (DMS and DLS) compared to the NAc in both metrics of release site density (p<0.05) and ΔF/F_0_ per site (p<0.05), with no significant differences observed between medial and lateral dorsal striatum (Figure 5a, b). A significant difference in the reuptake time constant was observed only between the DLS and NAc (p<0.05) (Figure 5c). Within the striatum of GDX males, regional variability was reduced. DMS and DLS metrics were more comparable to NAc (Figure 5), with the exception of difference in release site density between the DLS and NAc.

**Figure 5.**
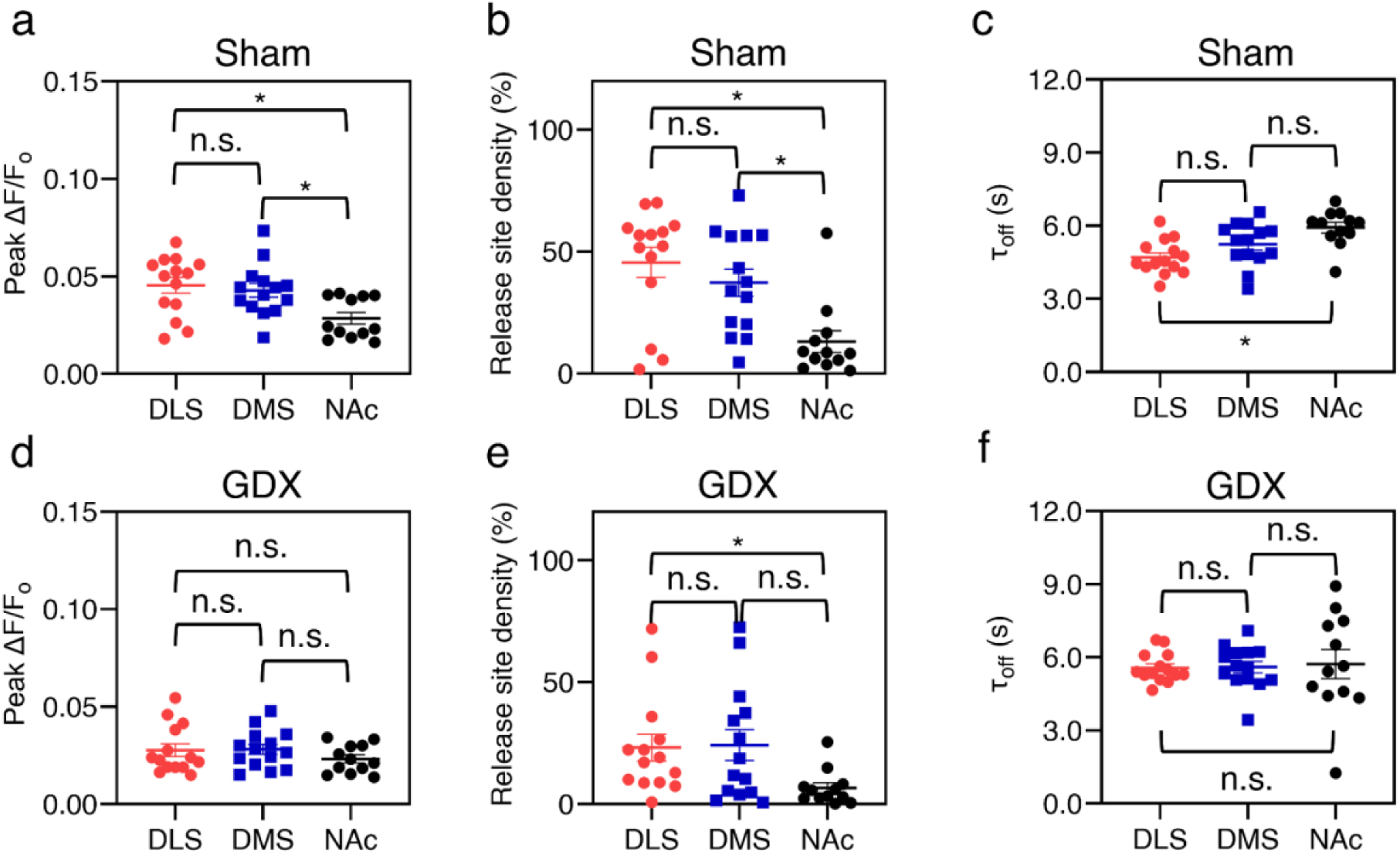
Prepubertal GDX eliminates regional differences in dorsal vs ventral evoked dopamine release, release site density, and reuptake. Comparison of metrics from sham (a-c) and GDX (d-f) mice. (a,d) Comparison of peak ΔF/F_0_ per release site between DMS, DLS, and NAc of (a) sham and (d) GDX mice shows dorsal vs. ventral differences in sham that are not apparent in GDX. (b,e) Comparison of release site density between the DMS, DLS, and NAc reveals dorsal vs. ventral differences in sham (b) that are only partially observed in GDX mice (e). (c,f) Comparison of τ_off_ between the DMS, DLS, and NAc reveals DLS vs NAc difference in sham (c) that is not observed in GDX mice (f). ANOVA with p = 0.05, followed by post-hoc analysis with Bonferroni correction. *p=0.05. Error bars represent the SEM.

## Discussion

We prepared acute brain slices from male *Mus spicilegus* mice raised on long day photoperiod to assess the influence of gonadal hormones at puberty on evoked dopamine release in the DMS, DLS, and NAc. We were interested in characterizing this species because its adolescent behaviors have been documented in the wild and its natural developmental programs are less likely to be disrupted by long-term laboratory breeding. We found that the absence of testes at puberty impacted evoked dopamine release in the nigrostriatal dopamine system, but not the mesoaccumbal system.

Using nIRCats enabled us to evaluate evoked dopamine release with spatial resolution divided into a grid of 7×7 µm ROIs. In the DLS and DMS, the peak of evoked nIRCat signals within active ROIs was significantly reduced in GDX mice when compared to sham controls. In the NAc, nIRCat signals in active ROIs were comparable between GDX and controls. In a spatial analysis of our grid of ROI data, we found that in the DLS, but not the DMS or NAc of GDX mice, the fraction of total ROIs that were active after stimulation was significantly smaller compared to sham controls. Finally, we found that in the DLS, but not the DMS or NAc of GDX mice, the τ_off_ was significantly longer compared to sham controls. These data support a working model in which a rise in gonadal hormones at puberty, likely testosterone or one of its metabolites, drives a gain in function in dopamine release from nigrostriatal terminals in the dorsal striatum, but does not impact mesoaccumbal terminal function. Further, this rise in pubertal hormones also drives an expansion in the density of active nigrostriatal terminals, as well as enhancing dopamine reuptake, in the DLS but not in other regions.

Our work has a number of limitations. We did not replace hormones in adulthood so we cannot distinguish between organizational or activational effects of gonadal steroids. We also did not study female mice so cannot say if puberty plays similar or different role in female development. Given the limitations of the single age sampled, we also cannot rule out the possibility of a situation in which complex responses to the GDX manipulation caused a diminishment in terminal release density and reuptake in the nigrostriatal system compared to sham controls. However, based on other literature, we favor the hypothesis that gonadal steroids in males at puberty drives enhancement in function of nigrostriatal terminals.

The published literature, largely using laboratory rats for experiments, establishes that the SNpc and VTA neurons that make dopamine are sensitive to gonadal steroids. Both estrogen and androgen receptors are expressed in nigrostriatal SNpc neurons in the midbrain (Quesada et al., 2007; Johnson et al., 2010; Kuhn et al., 2010; Purves-Tyson et al., 2012; Sárvári et al., 2014). The VTA mesoaccumbal system expresses estrogen receptors as well, but may have fewer androgen receptors and/or they may not be expressed in dopamine neurons (Kritzer, 1997). Testosterone from the testes could act on SNpc neurons by binding to androgen receptors, but it could also be converted to 17-beta estradiol through aromatization in the brain. However, in male rats, studies of adolescent GDX and comparison of GDX rats treated with testosterone, dihydrotestosterone, or estradiol suggest SNpc neurons can upregulate their TH and DAT mRNA expression in response to androgens but not estrogens during adolescence (Purves-Tyson et al., 2014). Interestingly, hormone-driven changes in TH and DAT were significant in soma samples of SNpc neurons but not significant in striatal tissue when sampling from their terminals (Purves-Tyson et al., 2014). This is not consistent with our data, but it is possible nIRCat imaging is simply a more sensitive metric to detect a change at terminals. Based on Purves Tyson et al. (2014), we favor a model in which activation of androgen receptors in the SNpc is the mechanism by which puberty supports trophic changes in DLS nigrostriatal dopamine release. This model can be falsified or supported in future experiments.

As we establish a mechanistic understanding of the development of the dopamine system by linking it to hormone receptor types and exploring changes at the level of terminal density, release and reuptake, we can better inform our understanding of developmental changes in exploratory behavior, motivation, salience, and learning during adolescence. This knowledge may provide mechanistic insights into understanding human adolescent vulnerability to anxiety and depression. Data from C57 Bl/6 mice show prepubertal GDX in male mice, but not female mice, reduces exploratory behavior in the elevated plus maze and enhances learned helplessness in the forced swim task (Boivin et al., 2017; Delevich et al., 2020). These are rodent tasks considered relevant to human anxiety and depression, and together with our current data suggest gonadal hormones at puberty play a prominent role in male mice in the development of bold and persistent behaviors, possibly through nigrostriatal dopamine.

Viewing these physiological and behavioral studies of prepubertal GDX from a neuroethological perspective may help us understand how pubertal processes could drive changes in behavior like dispersal, territory identification, and/or mate seeking (Lin and Wilbrecht, 2022). Our data are taken from *Mus spicilegus,* a wild-derived mouse known to disperse from their natal sites if they are born in the spring and delay their dispersal until the following spring if they are born in the winter (Garza et al., 1997; Gouat et al., 2003; Poteaux et al., 2008; Simeonovska-Nikolova and Mehmed, 2009; Cryns et al., 2022). Our mice were reared on 12:12h photoperiod in the lab which may mimic spring/summer light and to drive earlier maturation more in line with laboratory rodent life history. As their adolescent life history is seasonal, we are interested in investigating the development of the dopamine system in age-matched cohorts of *Mus spicilegus* reared on different photoperiods in future studies.

Our data are taken from *ex vivo* slice and so of course cannot fully capture functional changes that may be occurring across puberty *in vivo*. Neurons in the VTA and striatum have been shown to change their firing properties across adolescence. In anesthetized rats *in vivo*, VTA neurons were found to fire about 40% faster in adolescents than adults, have higher burst duration, greater number of spikes per burst, and shorter post-burst recovery period (McCutcheon et al., 2012). In awake-behaving rats, dorsal striatal neurons were, as a population, more active in adolescent rats than adults during a behavioral task, however, no difference in activity between age groups was observed in the ventral striatum (Sturman and Moghaddam, 2012). Further recording work *in vivo*, will be needed to examine how dopamine release changes with age and puberty in a task context and how this affects activity of specific downstream neurons. Developmental changes in dopamine in the nucleus accumbens shell and posterior tail of the striatum may also be worthy of investigation because these areas are also implicated in exploratory behavior (Nicola, 2010; Morrison et al., 2017; Menegas et al., 2018, Akiti et al., 2022; Tsutsui-Kimura et al., 2025).

In conclusion, our data show that dopamine terminal function, particularly in the dorsolateral striatum is altered by gonadal hormones at puberty in the male mouse brain. These data are only a piece of a larger puzzle, but they contribute to a growing body of evidence that the striatum does continue to develop during puberty (Larsen et al., 2020). This protracted maturation may have implications for striatal contribution to natural behaviors, which can be fruitfully explored in *Mus spicilegus*. These results may also help us to identify new mechanisms that play a potential role in the etiology of mental health issues in human adolescence.

## Acknowledgements

We acknowledge support of NIH R21DA059242 (LW), Burroughs Wellcome Fund Career Award at the Scientific Interface (CASI) (MPL), a Dreyfus foundation award (MPL), the Philomathia foundation (MPL), an NSF CAREER award 2046159 (MPL), a McKnight Foundation award (MPL), a Simons Foundation Award (MPL), a Moore Foundation Award (MPL), a Heising-Simons Fellowship (MPL), a Brain Foundation Award (MPL), and a polymaths award from Schmidt Sciences, LLC (MPL). MPL is a Chan Zuckerberg Biohub investigator, and a Helen Wills Neuroscience Institute Investigator. G.P., S.J., and J.M. contributed equally to this publication, and may rearrange the ordering of authorship when presenting this work.

## References

Abreu P, Hernandez G, Calzadilla CH, Alonso R (1988) Reproductive hormones control striatal tyrosine hydroxylase activity in the male rat. Neurosci Lett 95(1-3):213–7.

Akiti K, Tsutsui-Kimura I, Xie Y, Mathis A, Markowitz JE, Anyoha R, Datta SR, Mathis MW, Uchida N, Watabe-Uchida M (2022) Striatal dopamine explains novelty-induced behavioral dynamics and individual variability in threat prediction. Neuron 110(22):3789–3804.e9.

Bárdos B, Török HK, Nagy I (2024) Comparison of the exploratory behaviour of wild and laboratory mouse species. Behav Processes 217:105031.

Beyene AG, Delevich K, Del Bonis-O’Donnell JT, Piekarski DJ, Lin WC, Thomas AW, Yang SJ, Kosillo P, Yang D, Prounis GS, Wilbrecht L, Landry MP (2019) Imaging striatal dopamine release using a nongenetically encoded near infrared fluorescent catecholamine nanosensor. Sci Adv 5(7):eaaw3108.

Boivin JR, Piekarski DJ, Wahlberg JK, Wilbrecht L (2017) Age, sex, and gonadal hormones differently influence anxiety- and depression-related behavior during puberty in mice. Psychoneuroendocrinology 85:78–87.

Copeland WE, Worthman C, Shanahan L, Costello EJ, Angold A (2019) Early Pubertal Timing and Testosterone Associated With Higher Levels of Adolescent Depression in Girls. J Am Acad Child Adolesc Psychiatry 58(12):1197–1206.

Cryns NG, Lin WC, Motahari N, Krentzman OJ, Chen W, Prounis G, Wilbrecht L (2022) The maturation of exploratory behavior in adolescent Mus spicilegus on two photoperiods. Front Behav Neurosci 16:988033.

Dahl RE, Allen NB, Wilbrecht L, Suleiman AB (2018) Importance of investing in adolescence from a developmental science perspective. Nature 554(7693):441–450.

Deardorff J, Hayward C, Wilson KA, Bryson S, Hammer LD, Agras S (2007) Puberty and gender interact to predict social anxiety symptoms in early adolescence. J Adolesc Health 41(1):102–4.

Delevich K, Hall CD, Piekarski D, Zhang Y, Wilbrecht L (2020) Prepubertal gonadectomy reveals sex differences in approach-avoidance behavior in adult mice. Horm Behav 118:104641.

Delevich K, Klinger M, Okada NJ, Wilbrecht L (2021) Coming of age in the frontal cortex: The role of puberty in cortical maturation. Semin Cell Dev Biol 118:64–72.

Garza JC, Dallas J, Duryadi D, Gerasimov S, Croset H, Boursot P (1997) Social structure of the mound-building mouse Mus spicilegus revealed by genetic analysis with microsatellites. Mol Ecol 6(11):1009–17.

Gazzara RA, Fisher RS, Howard SG (1986) The ontogeny of amphetamine-induced dopamine release in the caudate-putamen of the rat. Brain Res 393(2):213–20.

Ge X, Conger RD, Elder GH Jr (2001) Pubertal transition, stressful life events, and the emergence of gender differences in adolescent depressive symptoms. Dev Psychol 37(3):404–17.

Gouat P, Féron C, Demouron S (2003) Seasonal reproduction and delayed sexual maturity in mound-building mice Mus spicilegus. Reprod Fertil Dev 15(3):187–95.

Graber JA, Seeley JR, Brooks-Gunn J, Lewinsohn PM (2004) Is pubertal timing associated with psychopathology in young adulthood. J Am Acad Child Adolesc Psychiatry 43(6):718–26.

Groó Z, Szenczi P, Bánszegi O, Altbäcker V (2013) Natal dispersal in two mice species with contrasting social systems. Behavioral Ecology and Sociobiology 67:235–242.

Hoops D, Flores C (2017) Making Dopamine Connections in Adolescence. Trends Neurosci 40(12):709–719.

Johnson ML, Day AE, Ho CC, Walker QD, Francis R, Kuhn CM (2010) Androgen decreases dopamine neurone survival in rat midbrain. J Neuroendocrinol 22(4):238–47.

Kritzer MF (1997) Selective colocalization of immunoreactivity for intracellular gonadal hormone receptors and tyrosine hydroxylase in the ventral tegmental area, substantia nigra, and retrorubral fields in the rat. J Comp Neurol 379(2):247–60.

Kuhn C, Johnson M, Thomae A, Luo B, Simon SA, Zhou G, Walker QD (2010) The emergence of gonadal hormone influences on dopaminergic function during puberty. Horm Behav 58(1):122–37.

Lafaille M, Féron C (2014) U-shaped relationship between ageing and risk-taking behaviour in a wild-type rodent. Animal Behaviour 97:45–52.

Larsen B, Olafsson V, Calabro F, Laymon C, Tervo-Clemmens B, Campbell E, Minhas D, Montez D, Price J, Luna B (2020) Maturation of the human striatal dopamine system revealed by PET and quantitative MRI. Nat Commun 11(1):846.

Lin WC, Wilbrecht L (2022) Making sense of strengths and weaknesses observed in adolescent laboratory rodents. Curr Opin Psychol 45:101297.

McCutcheon JE, Conrad KL, Carr SB, Ford KA, McGehee DS, Marinelli M (2012) Dopamine neurons in the ventral tegmental area fire faster in adolescent rats than in adults. J Neurophysiol 108(6):1620–30.

Mendle J, Ryan RM, McKone KMP (2018) Age at Menarche, Depression, and Antisocial Behavior in Adulthood. Pediatrics 141(1):e20171703.

Menegas W, Akiti K, Amo R, Uchida N, Watabe-Uchida M (2018) Dopamine neurons projecting to the posterior striatum reinforce avoidance of threatening stimuli. Nat Neurosci 21(10):1421–1430.

Morrison SE, McGinty VB, du Hoffmann J, Nicola SM (2017) Limbic-motor integration by neural excitations and inhibitions in the nucleus accumbens. J Neurophysiol 118(5):2549–2567.

Natterson-Horowitz B, Bowers K (2020) Wildhood: the astounding connections between human and animal adolescents. New York, NY: Scribner.

Nicola SM (2010) The flexible approach hypothesis: unification of effort and cue-responding hypotheses for the role of nucleus accumbens dopamine in the activation of reward-seeking behavior. J Neurosci 30(49):16585–16600.

Piekarski DJ, Johnson CM, Boivin JR, Thomas AW, Lin WC, Delevich K, M Galarce E, Wilbrecht L (2017) Does puberty mark a transition in sensitive periods for plasticity in the associative neocortex? Brain Res 1654(Pt B):123–144.

Pitts EG, Stowe TA, Christensen BA, Ferris MJ (2020) Comparing dopamine release, uptake, and D2 autoreceptor function across the ventromedial to dorsolateral striatum in adolescent and adult male and female rats. Neuropharmacology 175:108163.

Poteaux C, Busquet N, Gouat P, Katona K, Baudoin C (2008) Socio-genetic structure of mound-building mice, Mus spicilegus, in autumn and early spring. Biological Journal of the Linnean Society 93(4), 689–699.

Purves-Tyson TD, Handelsman DJ, Double KL, Owens SJ, Bustamante S, Weickert CS (2012) Testosterone regulation of sex steroid-related mRNAs and dopamine-related mRNAs in adolescent male rat substantia nigra. BMC Neurosci 13:95.

Purves-Tyson TD, Owens SJ, Double KL, Desai R, Handelsman DJ, Weickert CS (2014) Testosterone induces molecular changes in dopamine signaling pathway molecules in the adolescent male rat nigrostriatal pathway. PLoS One 9(3):e91151.

Quesada A, Romeo HE, Micevych P (2007) Distribution and localization patterns of estrogen receptor-beta and insulin-like growth factor-1 receptors in neurons and glial cells of the female rat substantia nigra: localization of ERbeta and IGF-1R in substantia nigra. J Comp Neurol 503(1):198–208.

Sárvári M, Deli L, Kocsis P, Márk L, Maász G, Hrabovszky E, Kalló I, Gajári D, Vastagh C, Sümegi B, Tihanyi K, Liposits Z (2014) Estradiol and isotype-selective estrogen receptor agonists modulate the mesocortical dopaminergic system in gonadectomized female rats. Brain Res 1583:1–11.

Shemisa K, Kunnathur V, Liu B, Salvaterra TJ, Dluzen DE (2006) Testosterone modulation of striatal dopamine output in orchidectomized mice. Synapse 60(5):347–53.

Simeonovska-Nikolova D, Mehmed S (2009) Behavior of mound-building mouse, Mus spicilegus during autumn-winter period in captivity. Biotechnology & Biotechnological Equipment 23(sup1):180–183.

Simpson EH, Gallo EF, Balsam PD, Javitch JA, Kellendonk C (2022) How changes in dopamine D2 receptor levels alter striatal circuit function and motivation. Mol Psychiatry 27(1):436–444.

Stamford JA (1989) Development and ageing of the rat nigrostriatal dopamine system studied with fast cyclic voltammetry. J Neurochem 52(5):1582–9.

Sturman DA, Moghaddam B (2012) Striatum processes reward differently in adolescents versus adults. Proc Natl Acad Sci U S A 109(5):1719–24.

Tsutsui-Kimura I, Tian ZM, Amo R, Zhuo Y, Li Y, Campbell MG, Uchida N, Watabe-Uchida M (2025) Dopamine in the tail of the striatum facilitates avoidance in threat-reward conflicts. Nat Neurosci 28(4):795–810.

Worthman CM, Trang K (2018) Dynamics of body time, social time and life history at adolescence. Nature 554(7693):451–457.

Xiao L, Becker JB (1994) Quantitative microdialysis determination of extracellular striatal dopamine concentration in male and female rats: effects of estrous cycle and gonadectomy. Neurosci Lett 180(2):155–8.

Yang SJ, Del Bonis-O’Donnell JT, Beyene AG, Landry MP (2021) Near-infrared catecholamine nanosensors for high spatiotemporal dopamine imaging. Nat Protoc 16(6):3026–3048.

Yin HH, Knowlton BJ (2006) The role of the basal ganglia in habit formation. Nat Rev Neurosci 7(6):464–76.

